# Lactoylglutathione promotes inflammatory signaling in macrophages

**DOI:** 10.1101/2023.10.10.561739

**Authors:** Marissa N. Trujillo, Erin Q. Jennings, Emely A. Hoffman, Hao Zhang, Aiden M. Phoebe, Grace E. Mastin, Naoya Kitamura, Julie A Reisz, Emily Megill, Daniel Kantner, Mariola M. Marcinkiewicz, Shannon M. Twardy, Felicidad Lebario, Eli Chapman, Rebecca L. McCullough, Angelo D’Alessandro, Nathaniel W. Snyder, Darren A. Cusanovich, James J. Galligan

## Abstract

Chronic, systemic inflammation is a pathophysiological manifestation of metabolic disorders. Inflammatory signaling leads to elevated glycolytic flux and a metabolic shift towards aerobic glycolysis and lactate generation. This rise in lactate corresponds with increased generation of lactoylLys modifications on histones, mediating transcriptional responses to inflammatory stimuli. Lactoylation is also generated through a non-enzymatic S-to-N acyltransfer from the glyoxalase cycle intermediate, lactoylglutathione (LGSH). Here, we report a regulatory role for LGSH in inflammatory signaling. In the absence of the primary LGSH hydrolase, glyoxalase 2 (GLO2), RAW264.7 macrophages display significant elevations in LGSH, while demonstrating a potentiated inflammatory response when exposed to lipopolysaccharides, corresponding with a rise in histone lactoylation. Interestingly, our data demonstrate that lactoylation is associated with more compacted chromatin than acetylation in an unstimulated state, however, upon stimulation, regions of the genome associated with lactoylation become markedly more accessible. Lastly, we demonstrate a spontaneous S-to-S acyltransfer of lactate from LGSH to CoA, yielding lactoyl-CoA. This represents the first known mechanism for the generation of this metabolite. Collectively, these data suggest that LGSH, and not intracellular lactate, is a primary contributing factor facilitating the inflammatory response.

## Introduction

The biology of lactate and lactate-derived post-translational modifications (PTMs) has recently seen a rapid resurgence in interest, with defined roles in cell cycle regulation,^1^ energy and redox homeostasis,^2^ the immune response,^3^ and chromatin function.^3–6^ The bulk of intracellular lactate is generated via glycolysis (up to 90%), and thus reducing lactate production has been a focal point in biomedical research.^7^ Recently, lactate has been identified as a source for the histone PTM, histone lactoylation.^3^ These PTMs are significantly elevated as lactate concentrations rise and are proposed to be derived from a lactoyl-CoA-dependent process. Lys residues are also prone to lactoylation through a non-enzymatic S-to-N acyltransfer from the glyoxalase cycle intermediate, lactoylglutathione (LGSH).^8^ These different sources for lactoylation are independent processes and thus it is important to investigate whether the source of this PTM (non-enzymatic vs enzymatic) is relevant in the context of different disease states, primarily those being metabolic. The relative contribution of these processes towards the total pool of protein lactoylation, however, is unknown.

Glycolysis is a tightly controlled cellular process, and alterations in flux are necessary for homeostasis. Glycolytic flux is elevated in response to numerous endogenous and exogenous cellular stresses (e.g. cancer and inflammation), leading to a rise in intracellular concentrations of glycolytic metabolites.^9–11^ Methylglyoxal (MGO) is perhaps one of the most heavily studied metabolic by-products of glycolysis, and has implications in nearly every disease state.^12–17^ Although the bulk of MGO is generated from the triose sugars, dihydroxyacetone phosphate (DHAP) and glyceraldehyde-3-phosphate (GA3P), MGO is also produced from lipid and amino acid metabolism.^18^ To combat increasing concentrations of this “toxic” metabolite, all cells are equipped with the glyoxalase cycle, consisting of two enzymes, glyoxalase 1 (GLO1) and GLO2.^19^ Within the glyoxalase cycle, MGO is conjugated to glutathione (GSH), forming a hemithioacetal (HTA) intermediate (**Figure 1a**). GLO1 then isomerizes the HTA to lactoylglutathione (LGSH). Lastly, GLO2 hydrolyzes LGSH to generate D-lactate and reintroduce GSH back into the cycle (**Figure 1a**).^20^ We have previously shown that LGSH, the intermediate in this cycle, serves as an acyl donor to available Lys residues, yielding the PTM, lactoylLys, through a non-enzymatic S-to-N acyltransfer (**Figure 1a**).^8^ These PTMs are enriched on glycolytic enzymes, reducing glycolytic output and lactate production.^5,8^ Lactoylation was also found on histones, mediating responses to hypoxic and inflammatory stimuli.^3^ These Lys lactoylation PTMs were proposed to be generated through an enzyme-catalyzed process using lactate-derived lactoyl-CoA and a putative lactoyltransferase, p300;^3^ the metabolic source and machinery required for lactoyl-CoA generation, however, has not been defined. This gap in knowledge is due, in part, to the analytical challenge in quantifying lactoyl-CoA, which exists at concentrations >1000-fold less than acetyl-CoA.^21^

**Figure 1.**
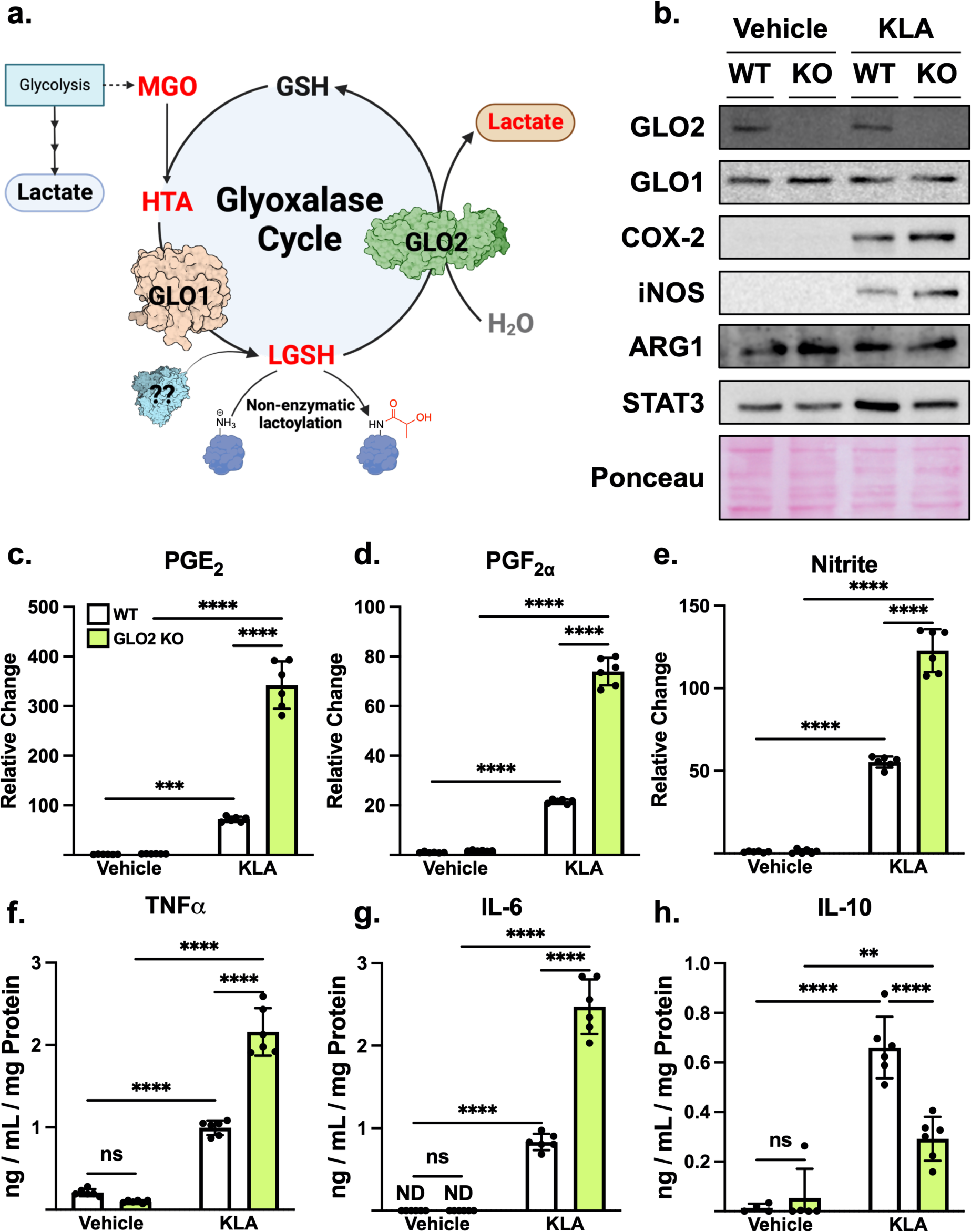
GLO2 ablation exacerbates the inflammatory response. **a.** The glyoxalase cycle detoxifies the glycolytic by-product, MGO, in a two-step process that cycles GSH. LGSH, the primary substrate for GLO2, serves as an acyl donor to free Lys residues yielding lactoylLys. **b.** KLA stimulation results in increased COX-2 and iNOS expression, which is exacerbated in KO cells. **c-e.** PGE_2_, PGF_2⍺_ and nitrite levels increase in KO cells following KLA treatment. **f-h.** Pro-inflammatory cytokines TNF⍺ and IL-6 are elevated in KO cells following KLA treatment while IL-10 (anti-inflammatory) is decreased. N=6 +/-SD. ****P < 0.0001, ***P < 0.001, **P < 0.01 by two-way ANOVA. Relative change: normalized to WT vehicle. Abbreviations: (L)GSH, (lactoyl)glutathione; MGO, methylglyoxal; HTA, hemithioacetal; GLO1, glyoxalase 1; GLO2, glyoxalase 2. KLA, Kdo_2_-Lipid A; COX-2, cyclooxygenase-2; iNOS, inducible nitric oxide synthase; ARG1, arginase 1; STAT3, signal transducer and activator of transcription 3; PG(E_2_, F_2⍺_), prostaglandin (E_2_, F_2⍺_); TNF⍺, tumor necrosis factor ⍺; IL(-6,10), interleukin(-6,10). *See also Figure S1*.

In this Research Article, we describe a regulatory role for GLO2 in response to inflammatory stimuli. GLO2 knockout RAW 264.7 macrophages display a significant increase in their response to the lipopolysaccharide mimetic, kdo_2_-Lipid A, as measured by prostaglandin and nitrite production, whose effects are independent of lactate production. We reveal a significant increase in LGSH production, also corresponding to significant increases in inflammatory mediators. Lastly, we demonstrate a potential metabolic source for lactoyl-CoA through a non-enzymatic S-to-S acyltransfer from LGSH to CoASH. Collectively, these data suggest that LGSH may serve as a major pro-inflammatory metabolite in macrophages.

## Results

### GLO2 suppresses the inflammatory response in RAW264.7 macrophages

Histone lactoylation has been proposed to promote adaptive responses to inflammatory stimuli and hypoxic conditions.^3^ As lactoylLys modifications are also generated through non-enzymatic means via S-to-N acyltransfer from the glyoxalase cycle intermediate, LGSH (**Figure 1a**),^8^ we sought to evaluate the contribution of GLO2 towards the inflammatory response using cultured RAW 264.7 macrophages. We generated GLO2 knockout (KO) cells and subjected them to a 24 h treatment with the chemically defined LPS mimetic, Kdo_2_-Lipid A (KLA),^22^ eliciting an inflammatory response similar to the conditions defined by Zhang et al. using LPS (**Figure S1**).^3^ As shown in **Figure 1b**, wild-type (WT) cells display a robust increase in the NFkB target proteins, cyclooxygenase-2 (COX-2) and inducible nitric oxide synthase (iNOS) following KLA treatment; this response is potentiated in KLA-challenged GLO2 KO cells, showing enhanced COX-2 and iNOS expression compared to WT cells. This is also corroborated through the measurement of COX-2-produced prostaglandin (PG) (**Figure 1c,d**) and iNOS-generated nitrite release in the culture medium (**Figure 1e**). WT cells show a significant increase in PG and nitrite production in response to KLA and these effects are significantly exacerbated in KO cells. We also quantified cytokine release, showing a significant increase in the pro-inflammatory cytokines, tumor necrosis factor α (TNFα) and interleukin 6 (IL-6) in WT cells in response to KLA with a further potentiation in KO cells (**Figure 1f-g**). Lastly, KO cells display blunted release of the anti-inflammatory cytokine, IL-10, compared to WT counterparts (**Figure 1h**). Collectively, these data suggest that GLO2 may regulate inflammatory signaling and provide us with the ability to investigate the potential role of non-enzymatically generated lactoylLys modifications in cultured macrophages.

### A putative metabolic source for lactoyl-CoA

Glycolysis-derived lactate is hypothesized to be the primary source for lactoyl-CoA in eukaryotic cells, serving as an acyl-donor for p300-catalyzed histone lactoylation (**Figure 2a**).^3,23^ We thus performed stable isotope tracing with ^13^C_6_-glucose in WT and KO cells treated with KLA for 24 h to quantify the fate of glucose carbon through glycolysis in WT and KO cells. As MGO is also generated through triose phosphate degradation, we quantified these metabolites as well. **Figure 2b** reveals a marked increase in ^13^C-labeled triose sugars in response to KLA, indicating increased flux through this pivotal point in glycolysis, regardless of genotype. Intriguingly, labeling of the terminal products of glycolysis (pyruvate, **Figure 2c** and lactate, **2d**), was unchanged in response to KLA, regardless of genotype. Our previous data has demonstrated that GLO2 KO HEK293 cells have a decrease in lactate production after MGO treatment and increased non-enzymatic lactoylLys generation.^8^ Thus, these data corroborate our findings and indicate that elevated glycolytic flux to lactate alone is insufficient to drive the increased inflammatory mediators observed in GLO2 KO cells.

**Figure 2.**
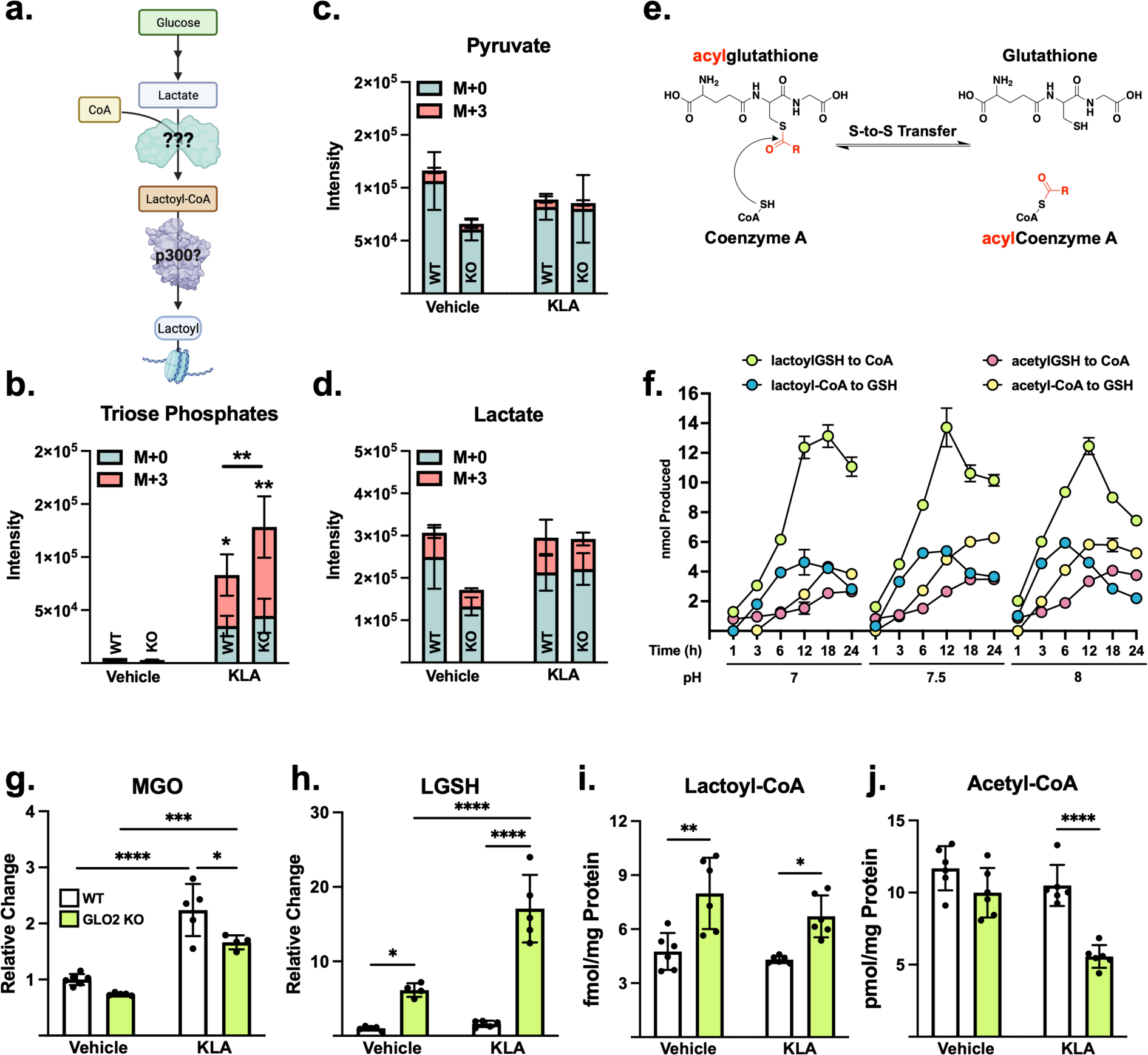
LGSH is a potential metabolic source for lactoyl-CoA. **a.** Lactoyl-CoA generation is proposed to be derived from lacate and CoA**. b-d**. Stable isotope labeling (SIL) of WT and KO cells treated with KLA reveals a significant increase in triose phosphates with no significant alterations in flux to pyruvate or lactate. **E.** The proposed mechanistic route for the generation of lactoyl-CoA from LGSH. **F.** Thioester migration is more efficient with lactate than acetate, indicating LGSH is a likely metabolic source for lactoyl-CoA. **g.** MGO is significantly increased in KLA treated cells, agreeing with **b. h.** LGSH is significantly increased in KO cells and this effect is potentiated in response to KLA. **i.** Lactoyl-CoA, and not **j.** acetyl-CoA, is increased in KO cells. N=6 +/-SD. ****P < 0.0001, ***P < 0.001, **P < 0.01, *P < 0.05 by two-way ANOVA. Relative change: normalized to WT vehicle. Abbreviations: CoA, Coenzyme A.

Lactoyl-CoA is present in eukaryotic cells at concentrations ∼120-350-fold lower than other acyl-CoA species.^21^ Although lactate is the presumed source of this metabolite, a mechanism for its esterification to CoA has not been established. Thioesters are heavily susceptible to nucleophilic attack by thiolate and amine-containing molecules, yielding a catalysis-free acyltransfer.^24^ This acyltransfer is heavily dependent on pH, favoring the acyl-transfer to CoA at lower pH as the pKa for the thiol GSH and CoA differ (8.4 and 9.8, respectively).^24^ This provides a pH-dependent balance of acyltransfer from acylGSH and/or acylCoA species in biological systems.^24,25^. We thus hypothesized that LGSH may serve as a metabolic source for lactoyl-CoA through this non-enzymatic mechanism, transferring the lactate moiety to CoASH (**Figure 2e**). To test this, we incubated equimolar concentrations of acylGSH or acylCoA analogs with their free thiol counterparts over a range of pH and time and quantified this acyltransfer using multiple reaction monitoring-mass spectromertry (MRM-MS). As shown in **Figure 2f**, the lactoyl group from LGSH readily transfers to CoA in a time-dependent manner; conversely, the transfer from lactoyl-CoA to GSH was markedly less efficient. We also performed the same experiment using acetyl-thioesters, showing minimal transfer regardless of the thioester used and at a much lower efficiency. These data indicate that the formation of lactoyl-CoA is favored, likely due to the differences in pKa between the GSH and CoA species, with free GSH having increased stability.

We next sought to evaluate this in cells. As expected, WT and KO cells display a significant increase in intracellular MGO levels following KLA treatment, indicating increased flux through the glyoxalase cycle (**Figure 2g**), which is consistent with the tracing in **Figure 2b**. As shown in **Figure 2h**, KO cells have elevated LGSH compared to their WT counterparts, consistent with our previous report;^8^ in response to KLA, KO cells display a further elevation in LGSH in response to KLA, while no significant alterations were observed in WT cells, which is consistent with the reported efficiency of the glyoxalase cycle.^20^ The abundance of lactoyl-CoA has not been determined in macrophages, despite its presumed regulatory role in inflammatory signaling.^3,23^ For this reason, we quantified lactoyl-CoA in these cohorts, revealing a significant elevation in KO cells compared with WT cells, while no alterations were observed in either genotype after KLA stimulation (**Figure 2i**). This indidates that perhaps lactoyl-CoA-derived lactoylation is not the primary source for the increased inflammatory response we observe. We also quantified acetyl-CoA (**Figure 2j**). Intrestingly, we demonstrate a significant reduction in this metabolite in KO cells following KLA treatment. These data indicate that KO cells do not have a global increase in acetylCoA species, consistent with the *in vitro* experiments in **Figure 2f**. Additionally, this demonstrates that lactoyl-CoA and acetyl-CoA generation may be inversely regulated, which ultimately may alter histone acylation. Furthermore, it should be noted that this is the first report of lactoyl-CoA quantitation in macrophages. Collectively, these data indicate that LGSH is capable of serving as an acyl donor and metabolic source for lactoyl-CoA in cells.

### Histone lactoylation is increased in KO cells in a site-specific manner

As histone lactoylation has been implicated in the inflammatory response, we next sought to quantify histone acylation marks. As observed through immunoblotting, we demonstrate a slight reduction in lactoylation of H3K18 upon stimulation with KLA, regardless of genotype (**Figure 3a**). In contrast, global histone lactoylation appears to be increased in KO cells, regardless of treatment. Akin to H3K18lac, basal histone acetylation is increased in the KO cells, whereas there is a decrease following KLA stimulation, regardless of genotype. As immunoblotting does not provide information on the relative abundance of PTMs compared to one another, we developed an MRM-MS assay to survey the histone PTM landscape, quantifying site-specific marks for Lys acetylation, Lys lactoylation, meLys, me_2_Lys, and me_3_Lys (**Figure 3b and S2**). Using this quantitative approach, we observed significant elevations in lactoylation in a site-specific manner, with H3K14, H3K18, H3.1K27, and H3K79 all significantly elevated compared to unstimulated cells (**Figure 3c,d,f,g**). The effects of GLO2 ablation, however, appear to be site-specific as only H3K79 and H3K8 are significantly elevated compared to WT counterparts (**Figure 3f,j**). Consistent with previous reports, we also observed a concomitant reduction in acetylation at these marks, suggesting stearic blockade by lactoylation.^26^ H3K27 is heavily investigated as me_3_Lys is highly associated with transcriptionally silenced regions of the genome while acetylation is considered a mark of active genomic regions.^27^ We note that acylation (acetylation and lactoylation) of this site is considerably low in abundance (< 1%) compared to methyl marks (**Figure 3l**). Overall, we observe a tight regulation of the lactoylation status of histones after KLA treatment in GLO2 KO cells, rather than a global elevation through the genome. Taken together, this demonstrates that acetylation and lactoylation are likely competing at multiple loci in a concerted, controlled manner, rather than a broad increase across the genome.

**Figure 3.**
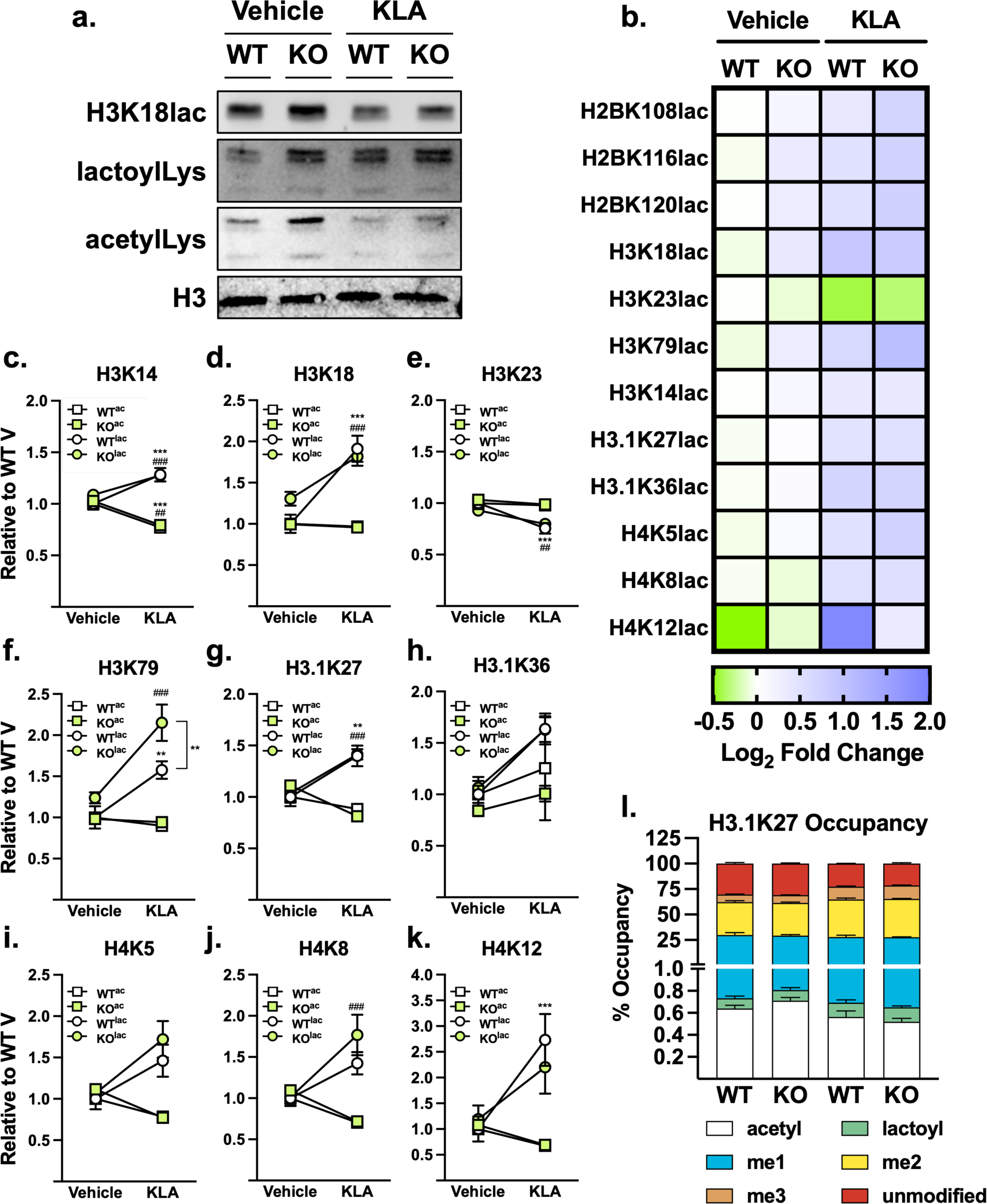
Histone lactoylation is elevated in a site-specific manner. **a.** Basal histone lactoylation is higher in KO cells, observed via immunoblotting. Representative blot, N=3. **b.** Specific histone sites have quantitative differences in lactoylation status in WT and GLO2 KO cells after inflammatory stimuli. **c-k.** Acetylation and lactoylation are dynamically regulated at specific loci, normalized to WT vehicle. **l.** Lactoylation is a low abundance mark compared to other canonical modifications. N=6 +/-SD. ***^/###^P < 0.001, **^/##^P < 0.01 by two-way ANOVA. *: WT to WT compared, ^#^: WT to KO compared. Abbreviations: ac, acetyl; lac, lactoyl; me, methyl. *See also Figure S2, Table S6*.

### Increased histone lactoylation leads to changes in chromatin organization consistent with gene activation at specific genomic loci

To explore the impact of GLO2 on chromatin accessibility dynamics at specific loci in the genome, we performed ATAC-seq on WT and KO cells following exposure to vehicle or KLA. In the vehicle-treated samples, we identified 110,907 peaks of accessibility with most (75.6%) common to the two genotypes (6,027 unique to WT, 20,988 unique to KO, and 83,892 common to both, **Figure 4a, Table S1**). For the KLA-stimulated samples, we identified a total of 126,553 peaks of accessibility, again with most (74.6%) common to the two genotypes (21,056 unique to WT, 11,038 unique to KO, 94,459 common to both, **Figure 4a, Table S1**).

**Figure 4.**
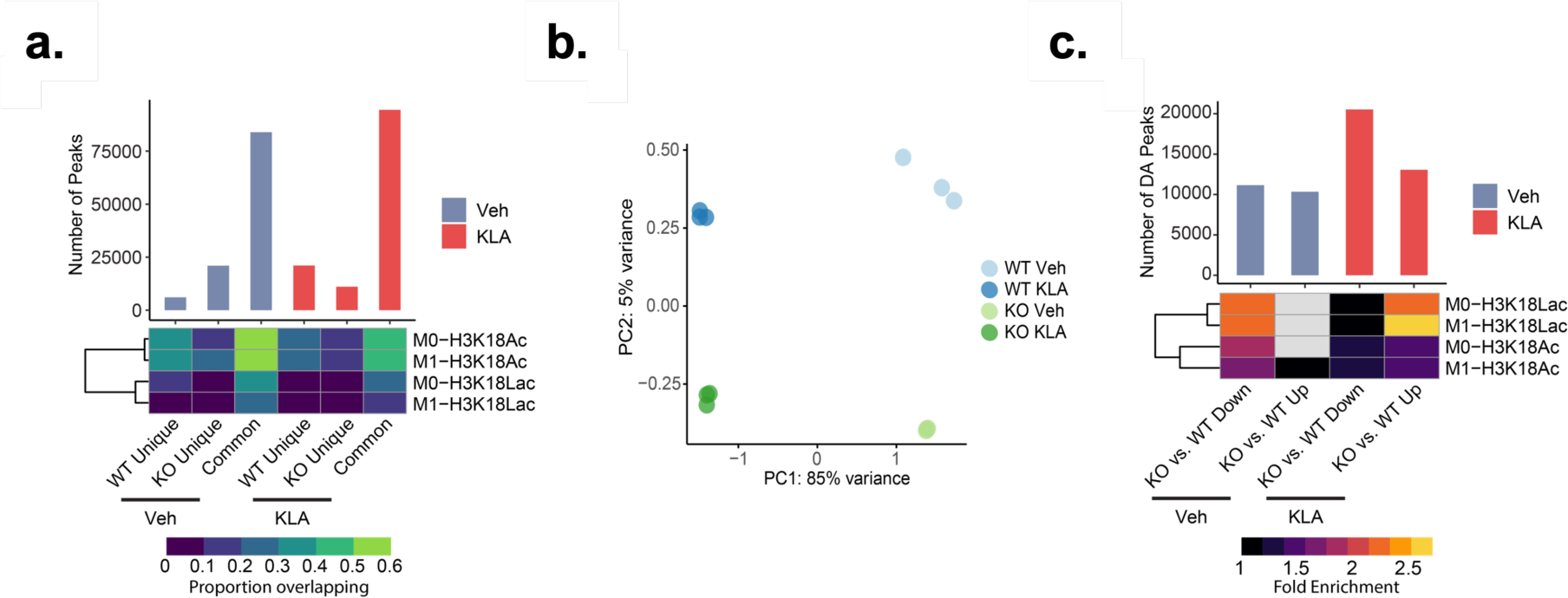
Differences in chromatin accessibility are attributed to increases in site specificic elevations of histone lactoylation. **a.** The number of peaks identified as either unique or common between WT and KO vehicle or between WT and KO KLA samples are shown. The heatmap visualizes the proportion of ATAC peaks overlapping with each ChIP-seq dataset. **b.** PCA analysis of ATAC-seq data where N=3. **c.** GLO2 KO-driven changes in chromatin accessibility are highly enriched in H3K18lac loci. Shown here are the number of differentially accessible peaks either ganining (Up) or losing accessibility (Down) in KO samples relative to the appropriate WT condition. Also indicated are the hypergeometric test enrichments for overlap of differentially accessible peaks identified in each contrast (column) with ChIP-seq datasets targeting H3K18ac and H3K18lac modifications in vehicle or LPS-treated macrophages (row, GEO accession: GSE115354). Only the tests having adjusted *p-values* lower than 0.05 are shown. *See also Figure S3, Table S1-S5*.

To better understand the role of histone lactoylation in chromatin accessibility in the KO samples, we intersected our data with previously published H3K18ac and H3K18lac ChIP-seq data generated in unstimulated and LPS-stimulated mouse bone marrow-derived macrophages (BMDMs).^3^ We found that both the H3K18ac and H3K18lac signals were most prevalent among common peaks (**Figure 4a**). Next, we sought to determine quantitative changes in accessibility in relation to genotype and stimulation. Thus, we combined all ATAC-seq peaks identified and quantified the relative accessibility of each peak (**Table S2**). Principal component analysis (PCA) and hierarchical clustering of the data indicate stratification of the samples first based on treatment and then by genotype (**Figure 4b and S3a**). While the genotype explained only 5% of the total variance present in the data, differential accessibility analysis revealed significant, widespread changes in chromatin accessibility due to the absence of GLO2 (**Figure 4c**). In untreated samples, we observed 10,343 loci with increased accessibility and 11,149 loci with decreased accessibility in the KO samples relative to WT; this difference is exacerbated following KLA treatment with 13,055 loci with increased accessibility and 20,527 loci with decreased accessibility. As H3K18lac is increased in both WT and KO cells (**Figure 3b,d**), we evaluated if the changes in chromatin accessibility were associated with these marks, specifically in the KO cells. To this end, we performed enrichment analysis on the differentially accessible sites identified in each group for overlap with the ChIP-seq data reported for H3K18lac ^3^. Interestingly, the differentially accessible sites identified between KO and WT cells (after treatment with either KLA or vehicle) were highly enriched in ChIP-seq peaks of H3K18lac (**Figure 4c**), suggesting that the altered chromatin accessibility caused by KO is likely attributable (at least partially) to the increase in H3K18lac.

We next examined whether the enrichments were generally associated with increasing or decreasing accessibility. In the KLA-stimulated samples, the enrichment of H3K18lac was primarily observed in sites that were more accessible in KO cells than WT. Strikingly, for the vehicle-treated cells, we observed the exact opposite trend, with enrichment of H3K18lac occurring primarily at sites that were less accessible in the KO cells. We interpret this to suggest that H3K18lac at baseline may be associated with less accessibility than H3K18ac; however, following stimulation, a yet-to-be-identified lactoylation reader may bind at sites associated with this modification and facilitate opening of the chromatin. To further explore the functional consequences of GLO2 KO in response to stimulation, we defined three major classes of dynamic loci – genomic loci exhibiting a consistent response to KLA treatment (regardless of genotype, N = 68,142), genomic loci uniquely responsive to the KLA treatment in WT cells (N = 16,712), and genomic loci uniquely responsive to KLA treatment in KO cells (N = 11,939) – and identified the pathways enriched among genes at these loci (**Table S3 and S4**). The common response pathways consistent with the expected response to KLA-stimulation include “antigen processing” and “apoptosis”, among others (**Figure S3b**). WT-specific responses were enriched for “alpha-linoleic acid metabolism,” “cGMP-PKG signaling,” and “fatty acid degradation,” among others (**Figure S3c**). On the other hand, KO-specific responses were characterized by the “Hippo signaling” pathway, the “TGF−beta signaling” pathway, and “porphyrin metabolism,” among others (**Figure S3c**). Collectively, these data indicate that the genomic responses observed in KO cells are largely augmented in response to KLA, consistent with the ChIP-seq data presented by Zhang et al.^3^ Furthermore, these results directly implicate an active reader of lactoylation on histone tails as a mediator of response to KLA stimulation.

### Non-enzymatic and enzymatic histone lactoylation are indistinguishable

The histone acetyltransferase, p300, has been proposed as an active histone lactoyltransferase, using lactoyl-CoA as a substrate.^3^ To explore epigenetic regulation of lactoylation in the context of GLO2, we treated WT and KO cells in the presence or absence of the p300 inhibitor, C646, a histone deacetylase inhibitor, trichostatin A (TSA), or the bromodomain (BET) inhibitor, JQ-1, with and without KLA stimulation (**Figure 5a, Figure S4**). Quantification of PGs in the presence of TSA reveals an exacerbation of the inflammatory response, particularly in KO cells (**Figure 5b,c, Figure S4**). This supports the proposed role of HDACs1-3 in the enzymatic removal of these PTMs on histones.^26^ Interestingly, treatment with C646 and KLA exacerbates PG production in KO cells relative to WT cells (**Figure 5b,c, Figure S4**). This further suggests that acetylation may serve as a stearic block for non-enzymatic lactoylation and the reduced enzymatic addition of acylation likely ‘frees up’ additional sites to undergo non-enzymatic modification.^3^ This also indicates that p300-mediated lactoylation is not major mechanism by which these marks are placed on histones. Lastly, JQ-1 appears to blunt these responses in both WT and KO backgrounds, further pointing to a putative bromodomain reader for these marks. Thus, the origins of these PTMs (enzymatic vs. non-enzymatic) are not overly distinguishable by reader domains (i.e. bromodomains), confirming the effects observed with TSA. All together, this data demonstrates that enzymatic regulators of histone acylation may also act on non-enzymatically derived PTMs.

**Figure 5.**
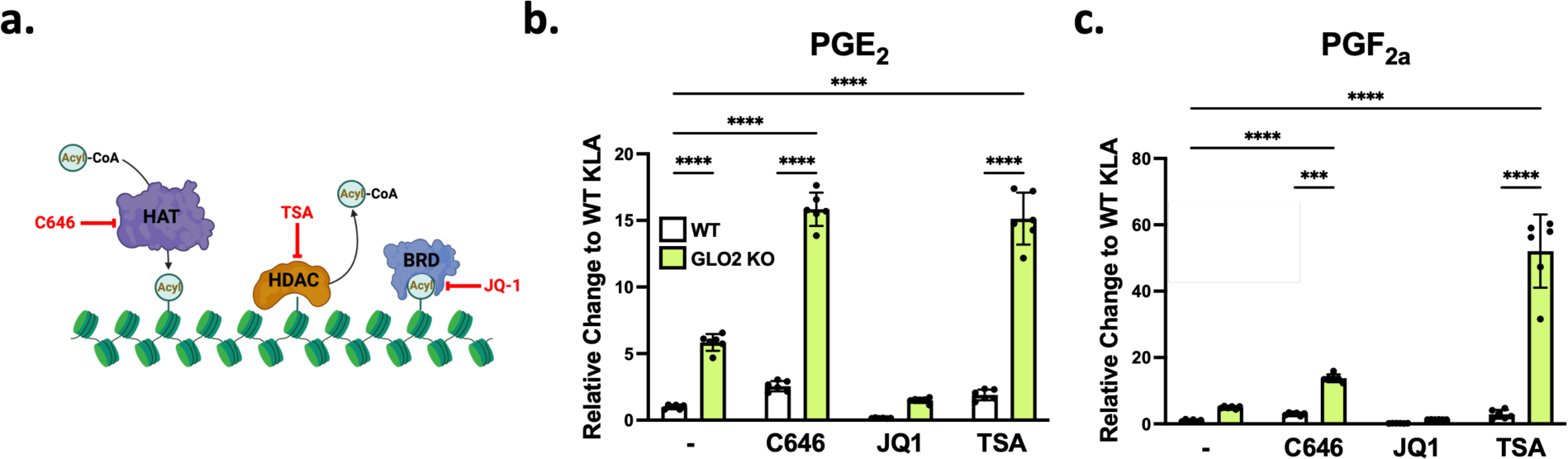
Enzymatic and non-enzymatic lactoylation are indistinguishable. **a.** Schematic representation of the inhibitor strategy used. **b,c.** PGs are significantly elevated in KO cells where HAT inhibition results in a significant increase in the inflammatory response. BET inhibition blunts this response regardless of the metabolic source of lactoylation. Lastly, HDAC inhibition results in a significant increase in PG production, indicating that both enzymatic and non-enzymatic marks are erased by HDACs on chromatin. Samples normalized to WT KLA-treated. Only KLA treated samples are shown. N=6 +/-SD. ****P < 0.0001, ***P < 0.001 by two-way ANOVA. Abbreviations: HAT, histone acetyltransferase; HDAC, histone deacetylase; BET, bromodomain and extra-terminal motif; TSA, trichostatin A. *See also Figure S4*.

## Discussion

In this Research Article, we describe a pro-inflammatory role for the glyoxalase cycle intermediate, LGSH, in macrophages and further, GLO2 regulation of the inflammatory response. LactoylLys has been shown to facilitate inflammatory responses in macrophages following exposure to LPS;^3^ these studies, however, lack mechanistic insight into how this PTM is generated in cells.^3^ We previously demonstrated that lactoylLys modifications can be generated through non-enzymatic means via the glyoxalase cycle intermediate, LGSH.^8^ As GLO2 is the primary enzyme responsible for controlling LGSH concentrations, cells lacking GLO2 have a marked elevation in LGSH and concomitant lactoylLys modifications.^8,19^ We thus explored the role of GLO2 in facilitating inflammatory signaling in cultured WT and GLO2 KO macrophages. Consistent with previous findings,^3^ we observe a significant increase in histone lactoylation upon KLA stimulation; these effects are elevated in a site-specific manner in GLO2 KO cells. Collectively, our data strongly supports a primary role for GLO2 as a driving factor in the inflammatory response.

Although lactoylLys has been proposed to be derived from lactoyl-CoA in an enzyme catalyzed reaction involving p300, this metabolite has not been quantified in macrophages, and its existence was only later found in cultured HepG2 cells.^3,21^ As S-to-S acyl transfers are well-established in biology,^24^ we hypothesized that lactoyl-CoA may be derived from LGSH, rather than through lactate and an unknown acyl-CoA synthetase. Indeed, this reaction takes place *in* vitro, with acyl transfer from LGSH to CoASH occurring more rapidly than its acetyl counterparts. Furthermore, the converse reaction (lactoyl-CoA to GSH) is not as favorable, perhaps suggesting that the equilibrium may be pushed to favor the generation of lactoyl-CoA *in vivo.* Further evidence for this mechanism being a major source for lactoyl-CoA generation in cells can be seen in **Figures 2i** and **j**, where an increase in lactoyl-CoA, and not acetyl-CoA, is observed in KO cells. In further support of this, no significant elevations in lactate production were observed in KO cells compared to WT counterparts using stable isotope labeling (**Figure 2d**). This result contradicts the proposed mechanism of lactoyl-CoA generation, which relies on esterification of lactic acid to CoASH via a yet-to-be-identified synthetase.^3^ This proposed mechanism has also been called into question by Dichtl and coworkers, who demonstrated that lactate (and thus lactate derived histone lactoylation) is a consequence, not a cause, of macrophage activation.^28^

Histone PTMs are often tightly regulated in a site-specific manner through enzyme-catalyzed reactions using so called “writers and erasers”.^29^ Current evidence has also supported that many PTMs may be derived through non-enzymatic means, with regulation occurring at the eraser-level.^25,30,31^ Indeed, we observe site-specific regulation of lactoylation, rather than broad changes across all identified sites. Zheng et. al demonstrated a significant increase in H3K18lac in response to LPS treatment.^3^ We show a similar trend in RAW264.7 cells treated with KLA. Competition with acetylation was also noted at this site, where increased lactoylation corresponded with a concomitant reduction in acetylation, indicating that this site is likely a critical signaling event for the inflammatory response. Indeed, bulk ATAC-seq results support this notion, where distinct sites within the genome were found to have altered accessibility, rather than broad changes across the genome. Furthermore, our ATAC-seq results indicate that lactoylation results in reduced accessibility at baseline relative to acetylation but dramatically increases accessibility upon stimulation, emphasizing the importance of the location of this mark and the consequent importance of maintaining an active eraser. Future work will be required to identify the implicated reader and additional factors regulating chromatin acessibility in this model. Treatment with C646, a p300 inhibitor, revealed a robust increase in PG production in GLO2 KO cells only. This is likely due to the increased availability of free Lys residues arising from reduced acetylation from HAT inhibition, allowing for additional non-enzymatic generation of histone lactoylation. This data also contradicts the proposed enzymatic generation of lactoylLys via p300. Lastly, our data indicates that the origins of lactoylLys are irrelvent to propagate inflammatory signaling as potent inhibition of PG production is still observed in KO cells treated with the BET inhibitor, JQ-1.

Overall, our data demonstrates that the glyoxalase cycle intermediate, LGSH, plays a regulatory role in inflammatory signaling. Of course, considerable work is still required to fully understand the role of LGSH, and thus GLO2, in the inflammatory response. LGSH seems to serve as a key metabolite that governs induction of numerous pro-inflammatory mediators, including cytokines, prostaglandins, and nitrite. Perhaps the most intriguing question is: *“what is the contribution of enzymatic vs. non-enzymatic lactoylation towards the inflammatory response?”* Although data presented in **Figure 5** attempts to address this question, the reduced occupancy of enzymatic marks stemming from pharmacological inhibition simply frees up additional marks to undergo non-enzymatic modification. This may indicate that at certain genomic loci, Lys acetylation may serve a protective role, shielding critical marks from undergoing non-enzymatic modification. A recent study by Mendoza et al. supports this notion, where Lys acetyl marks are removed from histone ‘resevoir sites’, recycled back into the acetyl-CoA pool, and then repositioned at ‘activating sites’.^32^ In this context, a similar mechanism may be taking place, where these ‘resevoir sites’ may confer protection against inappropriate lactoylation at ‘activating sites’. Fully unraveling this mechanism in cells, however, is a considerable analytical challenge due to the complexity and repurposing of cellular metabolites. Our model attempts to investigate the role of GLO2 (and elevations of LGSH) in macrophage function. We note the complex dynamics of immunity and that more work needs to be implemented to investigate the exact molecular mechanisms that are altered in the absence of GLO2. While ablation of GLO1 would seemingly provide some mechanistic insights into this regulation, we previously showed that GLO1 KO cells display no significant alterations in LGSH compared to WT counterparts, pointing to additional mechanisms of LGSH production.^8,33^ Future studies will aim to address understand how lactoyl-modified substrates are regulated both *in vitro* and *in vivo*.

## Acknowledgements

Financial support was provided by National Institutes of Health Grants: R35GM137910 and R01DK133196 for J.J.G.; T32 GM008804, for M.N.T. and N.K.; T32 ES007091 for E.Q.J; R35GM137896 for D.A.C; R01 CA259111, R01 GM132261 for N.W.S; R01 ES031463 for E.C.. Lactoyl-CoA was generously provided by Jordan Meier.

## Supplemental Information

**Figure S1. Related to Figure 1.**
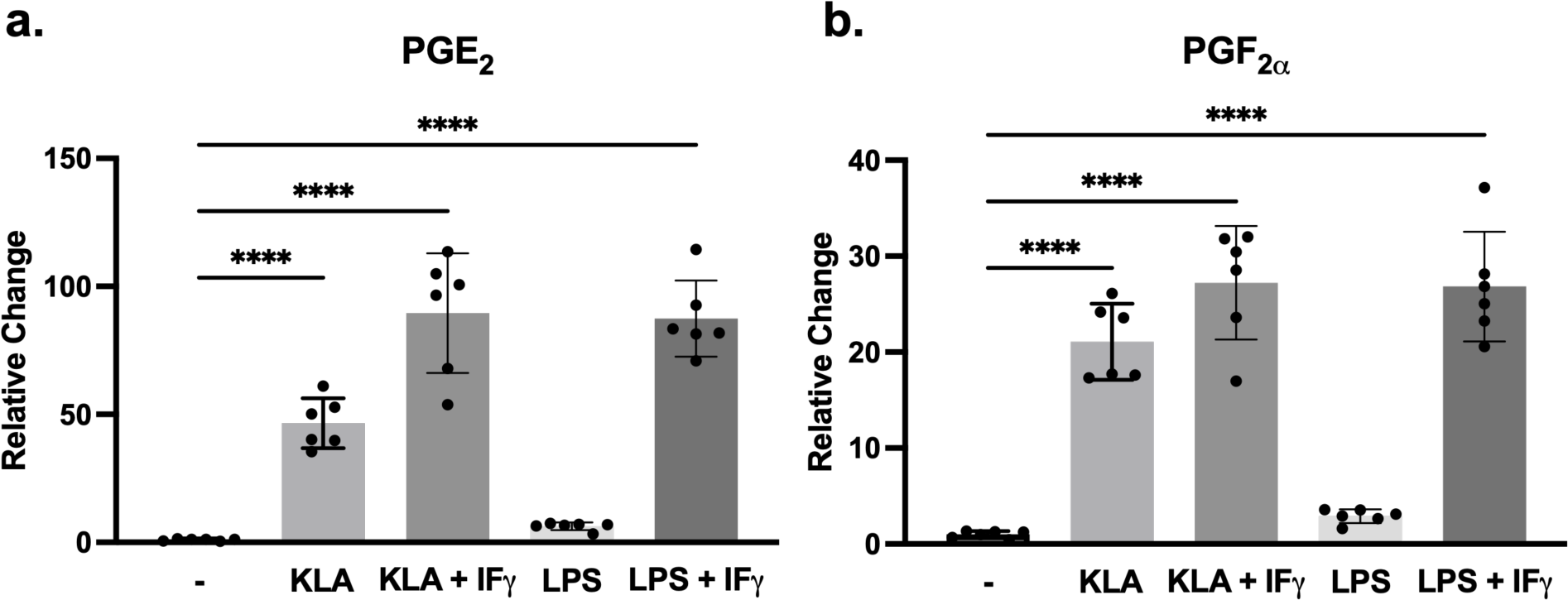
LPS + IFγ treatments and KLA treatments are comparable. **a-b**. RAW 264.7 macrophages treated with different inflammatory stimuli generate comparable levels of prostaglandins. N=6 +/-SD. ****P < 0.0001 by two-way ANOVA. Relative change: normalized to WT vehicle.

**Figure S2. Related to Figure 3.**
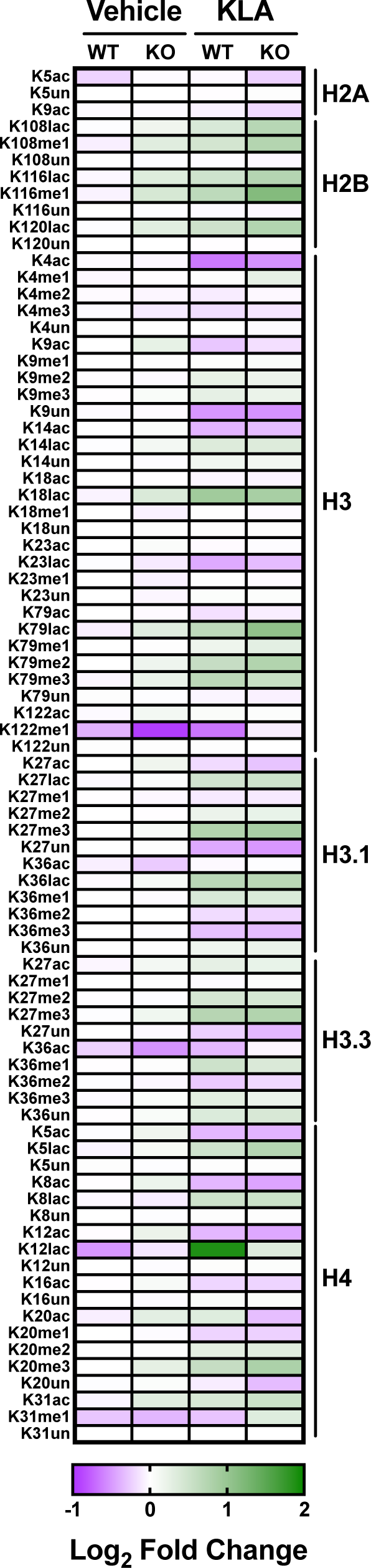
Quantification of histone post-translational modifications. Methylation (mono,di,tri), acetylation, lactoylation and unmodified histones sites quantified using MRM-MS. N=6 +/-SD. Abbreviations: me1,2,3, (mono, di, tri)methyl; ac, acetyl; lac, lactoyl.

**Figure S3. Related to Figure 4.**
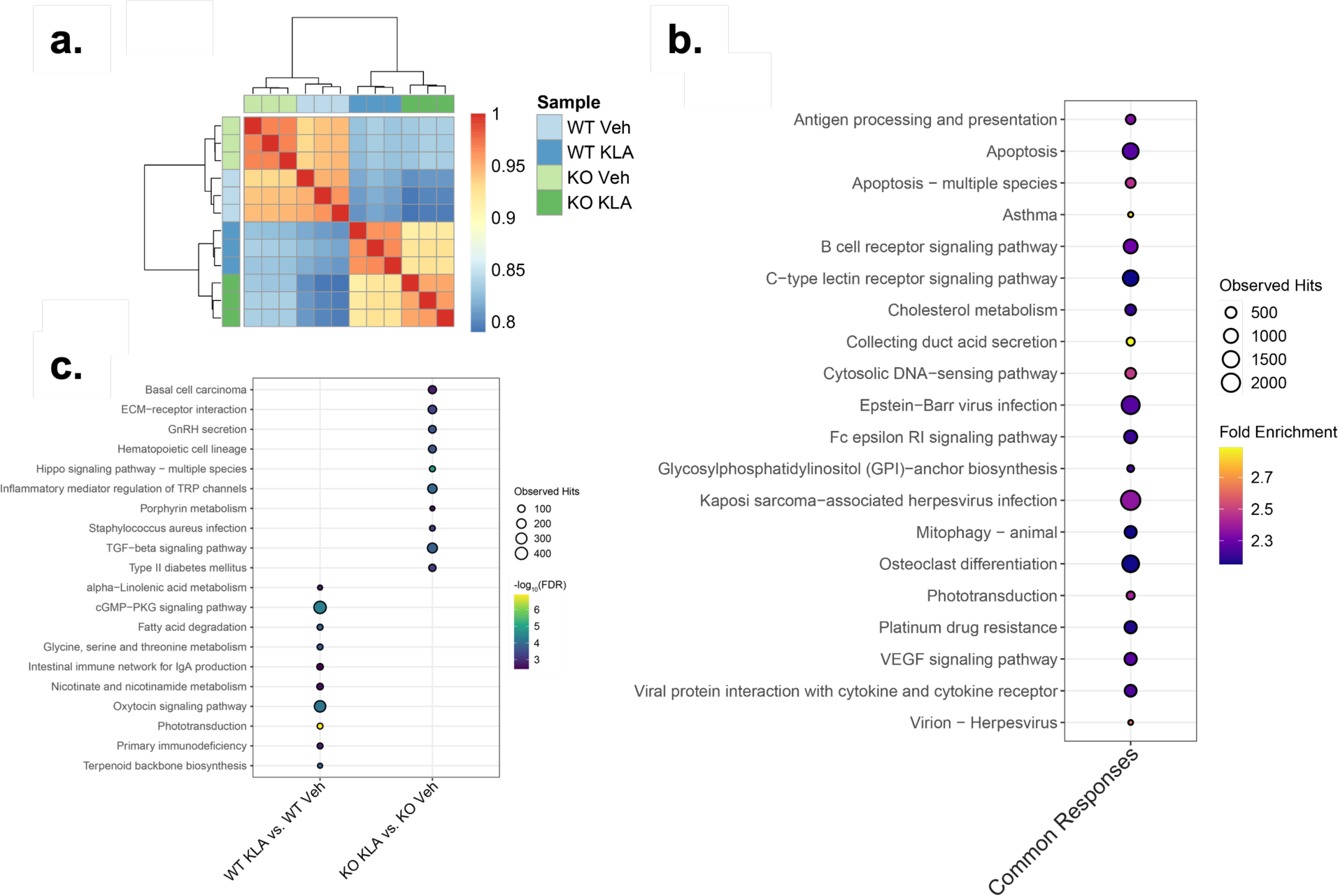
Functional analysis of dynamic regulatory elements induced by GLO2 KO and/or KLA treatment. **a.** Correlation heatmap of ATAC-seq data with three replicates in each group. The distance matrix was calculated as 1 - Pearson correlation coefficient. **b-d.** KEGG pathway enrichment in KLA-induced peaks either commonly (b) or uniquely (c) identified in WT and GLO2 KO samples. In panel (b), the top 20 enriched pathways with an adjusted *p-value* equal to zero are shown for common changes between genotypes in response to stimulation. The color code denotes the fold enrichment and dot size represents the number of regions observed in each pathway. The top 10 significant pathways that were uniquely identified in each contrast are shown in panel (c). The color represents the -log_10_-transformed adjusted *p-value* derived from the binomial tests. The dot size denotes the number of observed regions in each test.

**Figure S4. Related to Figure 5.**
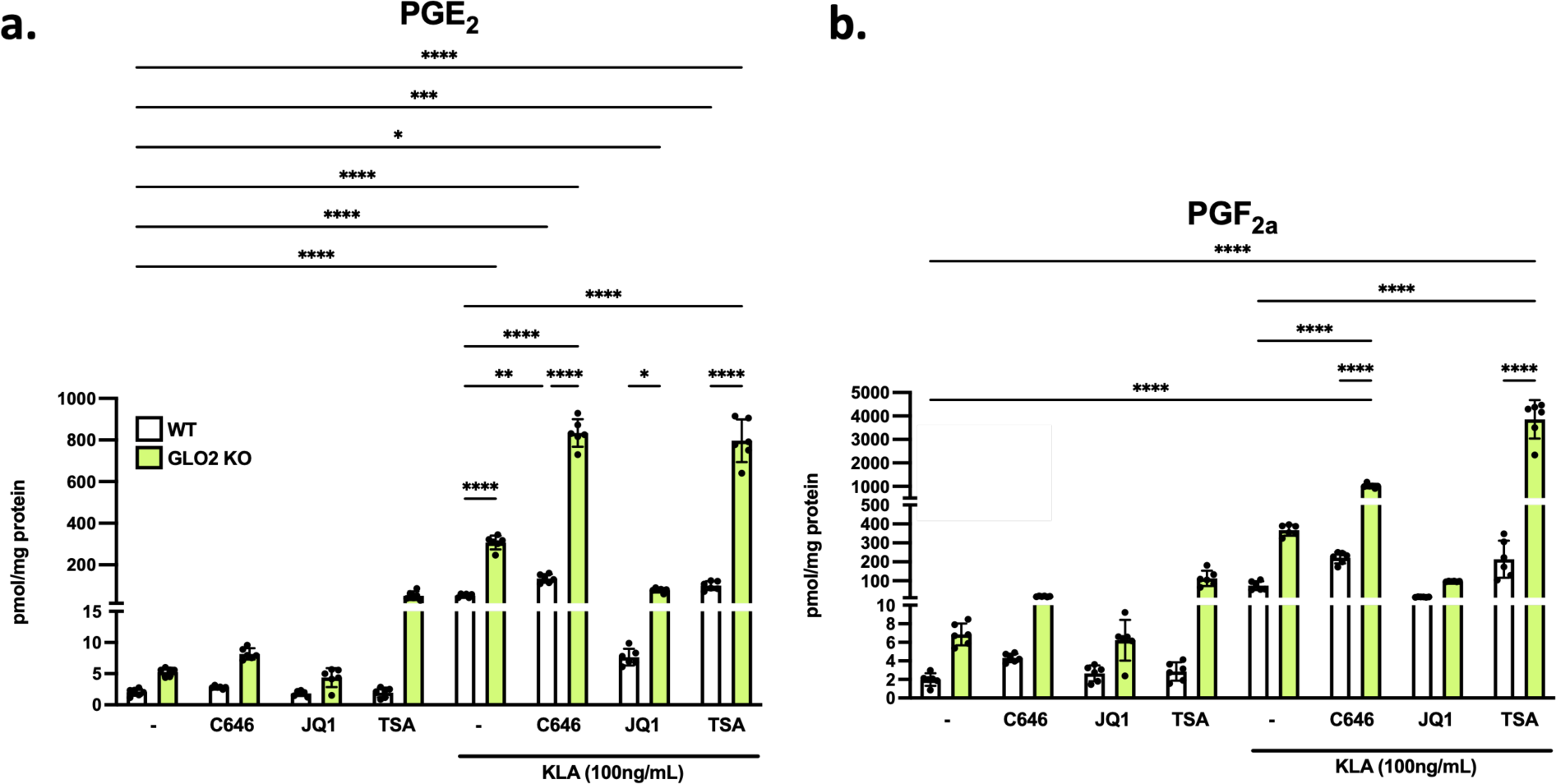
Prostaglandin production of WT and GLO2 KO macrophages treated with epigenetic inhibitors with and without inflammatory stimuli. **a,b.** PG production in WT and GLO2 KO macrophages that have been treated with three epigenetic inhibitors (Figure 5) with and without KLA treatment. N=6 +/-SD. ****P < 0.0001, ***P < 0.001, **P < 0.01, *P < 0.05 by two-way ANOVA.

**Table S1. Related to Figure 4.**

**Peaks of chromatin accessibility identified WT and GLO2 KO macrophages.** This table is provided as a separate file and contains the following columns:

Column 1 – “Chr”, chromosome name on which the peak was identified.

Column 2 – “Start”, start coordinate of peak.

Column 3 – “End”, end coordinate of peak.

Column 4 – “Genotype”, indicates if the peak was identified in WT or GLO2 KO cells.

Column 5 – “Treatment”, indicates if the peak was identified in vehicle or KLA treated cells.

**Table S2. Related to Figure 4.**

**Matrix of quantitative accessibility for ATAC-seq data.** This table is provided as a separate file and contains the following columns:

Column 1 – “peaks” for the chr_start_end identifier of each merged peak.

Columns 2-13 – each column indicates a genotype, a treatment, and a replicate ID (e.g. “glo2_kla1” is replicate 1 of the GLO2 KO, KLA-treated samples). Each cell of the matrix provides the number of reads for that sample overlapping the specified peak.

**Table S3. Related to Figure 4.**

**Pathway enrichment results for peaks with common responses in KLA treatment for WT and KO macrophages.** This table is provided as a separate file and contains the following columns:

Column 1 – “id” pathway term ID.

Column 2 – “genome_fraction”, fraction of genome covered by that term.

Column 3 – “observed_region_hits”, the number of peaks overlapping regions annotated as being associated with that pathway

Column 4 – “fold_enrichment”, observed/expected ratio. Column 5 – “p_value”, enrichment p-value.

Column 6 – “p_adjust”, multiple testing corrected p-value.

Column 7 – “mean_tss_dist”, average distance to nearest transcription start site for each peak.

Column 8 – “desc”, the textual description of the pathway.

**Table S4. Related to Figure 4.**

**Pathway enrichment results for peaks that respond uniquely to KLA treatment in WT and KO macrophages.** This table is provided as a separate file and contains the following columns:

Column 1 – “id”, pathway term ID.

Column 2 – “genome_fraction”, the fraction of genome covered by that term.

Column 3 – “observed_region_hits”, the number of peaks overlapping regions annotated as being associated with that pathway

Column 4 – “fold_enrichment”, observed/expected ratio. Column 5 – “p_value”, enrichment p-value.

Column 6 – “p_adjust”, multiple testing corrected p-value.

Column 7 – “mean_tss_dist”, average distance to nearest transcription start site for each peak.

Column 8 – “desc”, textual description of the pathway.

**Table S5. Related to Figure 4.**

**Sequencing barcodes used to demultiplex samples.** This table is provided as a separate file and contains the following columns:

Column 1 – “I7_Index_ID”, barcode ID.

Column 2 – “Index Sequence”, the actual sequence of the barcode used to demultiplex each sample.

Column 3 – “Genotype”, indicates “WT” or “Glo2KO”.

Column 4 – “Treatment”, indicates “Vehicle” or “KLA” treatment.

Column 5 – “Replicate”, indicates replicate number (1 to 3).

**Table S6. Related to Figure 3.**

**MRM-MS conditions for histone site quantifications.** This table is provided as a separate file and contains the parameters MS parameters for peptide identification and quantification.

## Materials and Methods

### Reagents

All reagents were purchased from ThermoFisher Scientific (Waltham, MA) unless otherwise stated. Fetal bovine serum (FBS) was purchased from Atlas Biologicals (Ft. Collins, CO). RAW 264.7 cells were purchased from ATCC.

### Cell culture

RAW264.7 cells were cultured in low-glucose DMEM supplemented with 10% FBS. Cells were incubated at 37°C under 5% CO_2_. Following treatments, cells washed and scraped into ice-cold PBS. Cells were pelleted via centrifugation at 1,000 × *g* for analysis.

### CRISPR-Cas9-mediated Glo2 knockout (^-/-^) RAW 264.7 cells

gRNA oligonucleotides were designed to target restriction enzyme recognition sites in exon 4 of the *GLO2* locus and ligated into the pSpCas9(BB)-2A-GFP plasmid according to Cong et al. ^1^. TATGGAGGTGATGACCGCAT was inserted into the plasmid to target GLO2. To generate G*LO2^-/-^*, 2 × 10^5^ RAW 264.7 cells were plated in 2 mL DMEM supplemented with 10% FBS in 6-well plates. The following day, 8 µg of each construct was combined with 10 µL lipofectamine 2000 (Life Technologies, Carlsbad, CA) reagent in 1 mL Opti-MEM and incubated at room temperature for 15 min. The DMEM was replaced with the plasmid-lipofectamine solution, and the cells were incubated at 37 °C for 24 h. The medium was then replaced, and cells were allowed to recover for 24 h at 37°C. Cells were then harvested for single cell sorting GFP positive cells. Colonies were verified for GLO2 knockout via immunoblotting.

### SDS-PAGE and immunoblotting

Samples were denatured in SDS loading buffer and heated at 95°C for 5 minutes. Proteins were then resolved via SDS-PAGE and transferred to nitrocellulose membranes (Biorad). Membranes were blocked with Odyssey Blotting Buffer (Li-Cor Biosciences, Lincoln, NE) for 45 minutes at room temperature. Primary antibodies were incubated with membranes overnight at 4°C as described: GLO1 (1:000, Protein Technologies, 15140), ARG1 (1,000, Gentex, GTX109242), STAT3 (1:1,000, Cell Signaling Technologies, 4904S), GLO2 (1:1000, ThermoFisher, PA5-28292), iNOS (1:2000, Cell Signaling Technologies, #2024), COX-2 (1:2000, Cell Signaling Technologies, #3582), H3K18La (1:1000, PTMBio, PTM-1406), LactoylLys (1:500, PTMBio, PTM-1401), AcetylLys (1:500, Abcam, Ab21623), H3 (1:20,000, Cell Signaling Technologies, #3638S), Actin (1:10,000, Sigma, #A1978). Following 3x washes with TBS +0.1% Tween-20 (TBST), infrared secondary antibodies (Li-COR) were added in blocking buffer (1:5,000) for 45 minutes. Blots were developed following 3 additional washes with TBST using a c600 Azure Imaging System (Azure Biosystems, Dublin, CA).

### Quantification of prostaglandins

5 × 10^5^ cells were plated in 6-well plates and allowed to seed overnight. Cells were then treated accordingly for 24h. For KLA treatments, cells were treated with either vehicle or KLA (100 ng/mL) for 24h. For LPS + IFγ treatments, cells were treated with LPS (5 ng/mL), IFγ (12 ng/mL), or the combination for 24h. For epigenetic inhibitor treatment, cells were treated with vehicle ± KLA (100 ng/mL), C646 (5µM) ± KLA (100 ng/mL), JQ1 (400 nM) ± KLA (100 ng/mL), or TSA (100 nM) ± KLA (100 ng/mL) for 24h. Following treatments, media (2 mL) for each well was removed and added to a tube of ethylacetate + 0.1% acetic acid (4 mL) containing 100 pmol PGE_2_-d4 and 100 pmol PGF2a-d_4_. Tubes were inverted 3X. Following separation, the supernatant (containing PGs) was removed and allowed to dry under N_2_. Samples were resuspended in MeOH (150 µL) and chromatographed (12 µL) using a Shimadzu LC system equipped with a 50 × 2.1mm, 3µm particle diameter Atlantis C_18_ column (Waters, Milford, MA) at a flow rate of 0.400 mL/min. Buffer A (0.1 formic acid (FA) in water) was held at 80% for 0.25 min then lowered to 1% over 5 min. Buffer B (0.1% FA in acetonitrile (ACN)) was held at 99% for 0.5 min and dropped to 20% B after another 0.5 min. The column was equilibrated at 20% B for 1.5 min. Multiple reaction monitoring (MRM) was conducted in negative mode using an AB SCIEX 6500 QTRAP+ with the transitions below. PGs were normalized to total protein (determined via BCA) per sample.

| Q1 mass | Q3 mass | ID | DP (volts) | CE (volts) |
| --- | --- | --- | --- | --- |
| 351.0 | 271.0 | PGE<sub>2</sub> | -60.0 | -23.0 |
| 355.0 | 275.0 | PGE<sub>2</sub>-d<sub>4</sub> | -60.0 | -23.0 |
| 353.2 | 193.1 | PGF<sub>2α</sub> | -120.0 | -34.0 |
| 357.2 | 197.1 | PGF<sub>2α</sub>-d<sub>4</sub> | -120.0 | -34.0 |

### Quantification of nitrite

5 × 10^5^ cells were plated as described above. Following treatments, media (1 mL) was reserved for nitrite quantification. Nitrite concentration was determined using the Griess assay^2^. Briefly, a standard nitrite curve was prepared using serial dilutions (100 µM, 50 µM, 25 µM, 12.5 µM, 6.25 µM, 3.125 µM, 1.56 µM, 0 µM nitrite). Sulfanilamide solution (1% sulfanilamide in 5% phosphoric acid) and NED solution (0.1% N-1-napthylethylene-diaminehydrochloride in water) were prepared ∼30 min prior (room temp) to assay measurements. 50 µL of media from treatments was removed and plated on a 96-well plate. Sulfanilamide solution (50 µL) was added and incubated for 10 min in the dark. NED solution (50 µL) was then added and incubated 10 min in the dark. Absorbance was then measured (520 nm) within 30 min. Nitrite was normalized to protein concentration (determined via BCA).

### Quantification of inflammatory cytokines

5 × 10^5^ cells were plated and treated as described above. Following treatment, media was removed for cytokine quantification and stored at -80°C until use. Cells were washed 1X with ice-cold PBS and scraped into ice-cold PBS. Samples were centrifuged (1,000 × *g*, 5 min, 4°C) and pellets were resuspended for total protein concentration, determined via BCA. Samples were normalized to total protein. Cytokine release was measured in cell culture supernatants using ELISA per the manufacturer’s protocol with slight modifications (Biolegend). Briefly, supernatants were thawed on ice and plated on coated ELISA plates (diluted 2-5 fold in diluent) with the appropriate freshly prepared standard and incubated overnight, rocking at 4°C. The following day, plates were washed 4x with 0.1% TBST and detection antibody was added for 1 h. Next, plates were washed with TBST an Avidin-HRP was added, and plates were incubated for 30 min in the dark. Finally, plates were washed with TBST, developed with TMB, and the reaction was stopped with 2M HCL. Development time varied (3-20 min) and the absorbance was measured at 450 nm.

### Quantification of cellular MGO

MGO was derivatized as previously described ^3^. Briefly, 5 × 10^5^ cells were plated in 6-well plated. The following day, cells were treated accordingly for 24h. Following treatment, media was removed, and plates were washed 1X and scraped into ice-cold PBS. After centrifugation (1,000 × *g*, 5 mins, 4°C), PBS was removed, and samples were resuspended in ice cold 80:20 MeOH: H_2_O (100 µL) containing 50 pmol ^13^C_3_-MGO. Protein was then precipitated for 1h at -80°C. Following centrifugation (14,000 × *g*, 10 mins, 4°C), supernatant was transferred to a tube containing 10 mM o-phenylenediamine (10 µL) and allowed to derivatize for 2h rotating end-over-end in the dark (RT). Samples were chromatographed (12 µL) using a Shimadzu LC40 system equipped with a 50 × 2.1mm, 3 µm particle diameter Atlantis C_18_ column (Waters, Milford, MA) at a flow rate of 0.500 mL/min. Buffer A (0.1% FA in water) was held at 95% for 0.5 min then a linear gradient to 98% buffer B (0.1% FA in ACN) was applied over the next 3 min. The column was held at 98% B for 2 min and then washed at 10% B for 0.5 min and equilibrated to 95% A for 2.5 min. Multiple reaction monitoring (MRM) was conducted in positive mode using an AB SCIEX 6500 QTRAP+ with the following transitions: *m/z* 145.0→77.0 (analyte); *m/z* 148.0→77.0 (^13^C_3_-MGO, internal standard). Samples were normalized to total protein concentration, determined via BCA.

### Quantification of GSH and LGSH

1 × 10^6^ cells were plated in 100 mm plates. The following day, cells were treated accordingly for 24h. Following treatment, media was removed, and plates were washed 1X and scraped into ice-cold PBS. Cells were pelleted via centrifugation at 1,000 × *g* (5 mins, 4°C), and resuspended in 20% 5-sulfosalcylic acid ((w/v) 150 µL) containing 1.25 nmol of GSH-(glycine-^13^C_2_,^15^N, internal standard). Samples were briefly sonicated and then centrifuged (14,000 × *g*, 10 mins). Supernatant (12 µL) was then chromatographed using a Shimadzu LC system equipped with a 150 × 3mm, 3 µm particle diameter Atlantis C_18_ column (Waters, Milford, MA) at a flow rate of 0.400 mL/min. Solvent A (10 mM heptafluorobutyric acid (HFBA) in H_2_O) was held at 95% for 1 min, and lowered to 90% A over the next 9 min. Then, a linear gradient to 98% B (10 mM HFBA in ACN) was applied over the next 10 min. The column was held at 98% B for 4.5 min and then equilibrated to 95% A for 0.5 min. The needle was washed prior to each injection with a buffer consisting of 25 mM NH_4_OAc in MeOH for 2.5 min. MRM was performed in positive ion mode using an AB SCIEX 6500+ QTRAP with the parameters below. GSH and LGSH were quantified using GSH-(glycine-^13^C_2_,^15^N). Samples were normalized to total protein.

| Q1 mass | Q3 mass | ID | DP (volts) | CE (volts) |
| --- | --- | --- | --- | --- |
| 308.0 | 179.0 | GSH | 51.0 | 17.0 |
| 311.0 | 182.0 | GSH-(glycine-$^{13}\text{C}_2$, $^{15}\text{N}$) | 51.0 | 17.0 |
| 380.1 | 233.1 | LGSH | 45.0 | 23.0 |

### Stable Isotope Labeling of RAW264.7 macrophages

5 × 10^5^ cells were plated and seeded overnight in 6-well plates. The following day, cells were treated with ^13^C_6_-glucose (4.375 mM) ± KLA (100 ng/mL) for 24h. Following treatments, cells were washed 3x with ice-cold PBS and scraped into 1 mL ice-cold PBS (2 wells were combined per treatment for an n = 3). Prior to centrifugation, a 10 µL aliquot was removed for cell counting. Samples were then centrifuged (1,000 × *g*, 5 min, 4°C) and supernatant was removed. Metabolites were extracted from frozen cell pellets at 4.0 × 10^6^ million cells per mL by vigorous vortexing in the presence of ice cold 5:3:2 MeOH:MeCN:water (v/v/v) for 30 min at 4°C. Following vortexing, supernatants were clarified by centrifugation (10 min, 12,000 × *g*, 4°C) and an aliquot of each extract was diluted 1:1 with the aforementioned solution during transfer to autosampler vials. The resulting samples were analyzed (10 µL per injection) by ultra-high-pressure liquid chromatography coupled to mass spectrometry (UHPLC-MS — Vanquish and Q Exactive, Thermo Fisher). Metabolites were resolved on a Kinetex C_18_ column using a 5 min gradient method exactly as previously described^4^. Following data acquisition, .raw files were converted to .mzXML using RawConverter then metabolites assigned and peaks integrated using Maven (Princeton University) in conjunction with the KEGG database and an in-house standard library. ^13^C Isotopic enrichment was visualized using GraphPad Prism 9.0. ^13^C_2_ metabolite peak areas were corrected for natural abundance. Quality control was assessed as using technical replicates run at beginning, end, and middle of each sequence as previously described^4^.

### Generation and quantification of acyl-CoAs and their glutathione counterparts

Generation of lactoyl-CoA, lactoylGSH, acetyl-CoA and acetylGSH were quantified using MRM-MS. Each reaction was incubated with equimolar CoA or GSH species in PBS at 37°C at the indicated pH and time. Reactions were quenched with 40% sulfosalicylic acid (SSA, 10 µL) and spiked with GSH-(glycine-^13^C_2_,^15^N) (GSH-IS). Samples were chromatographed (12 µL), using a Shimadzu LC system equipped with a 150 × 3mm, 3 µm particle diameter Atlantis C_18_ column (Waters, Milford, MA) at a flow rate of 0.450 mL/min. Solvent A (10 mM HFBA in H_2_O) was held at 92.5% for 1 min and lowered to 2% A over the next 3 min. The column was held at 98% buffer B (10 mM HFBA in ACN) for 2 mins. A was then increased to 92.5% for 0.5 min and equilibrated at 92.5% A for 2.5 min. The needle was washed prior to each injection with a buffer consisting of 25 mM NH_4_OAc in MeOH. MRM was performed in positive ion mode using an AB SCIEX 6500+ QTRAP with the parameters below. Analytes were quantified using a standard curve against GSH-IS.

| Q1 mass | Q3 mass | ID | DP (volts) | CE (volts) |
| --- | --- | --- | --- | --- |
| 308.0 | 179.0 | GSH | 51.0 | 17.0 |
| 311.0 | 182.0 | GSH-(glycine-$^{13}\text{C}_2$, $^{15}\text{N}$) | 51.0 | 17.0 |
| 350.0 | 203.0 | acetylGSH | 50.0 | 22.0 |
| 380.1 | 233.1 | LGSH | 45.0 | 23.0 |
| 810.0 | 303.0 | Acetyl-CoA 1 | 81.0 | 45.0 |
| 810.0 | 428.0 | Acetyl-CoA 2 | 81.0 | 45.0 |
| 768.0 | 261.0 | CoA 1 | 81.0 | 45.0 |
| 768.0 | 468.0 | CoA 2 | 81.0 | 45.0 |
| 840.5 | 333.0 | Lactoyl-CoA 1 | 85.0 | 43.0 |
| 840.5 | 428.0 | Lactoyl-CoA 2 | 120.0 | 40.0 |

### Acyl-CoA Quantification

2.5 × 10^6^ cells were plated and incubated overnight. Cells were treated accordingly for 24h. Following treatments, media was removed, and cells were washed 1X with ice-cold PBS. Cells were then scraped into 1 mL of ice-cold 10% trichloroacetic acid (TCA) and pelleted (1,000 × *g*, 5 min, 4°C). LC-MS/HRMS was performed as described^5^. Briefly, cell pellets and synthetically prepared acyl-CoA standards were resuspended in 1 mL of ice-cold 10% trichloroacetic acid (Sigma-Aldrich). Samples and calibrators were spiked before extraction with ^13^C_315_N_1_-acyl-CoAs made biosynthetically in HepG2 cells serially passaged in stable isotope labeling from essential nutrients in cell culture using ^13^C_315_N_1_-pantothenate. Samples were then sonicated for 10 × 0.5 second pulses to completely disrupt cellular membranes and incubated on ice to precipitate proteins. Protein was pelleted at 16,000 × RCF for 10 min at 4°C. Supernatant was collected and purified by solid-phase extraction using Oasis HLB 1cc (30 mg) SPE columns (Waters). Eluate was evaporated to dryness under N_2_ gas and re-suspended in 50 μL of 5% SSA (w/v) for injection. 20 μL of each sample was analyzed on a Vanquish Duo UHPLC with a Waters HSS T3 (2.1 × 100mm, 2.7 µm) column couple to a Q Exactive Plus (Thermo Scientific). A single ion monitoring (SIM) acquisition was used for lactoyl-CoA to increase sensitivity. The ratio of the peak for lactoyl-CoA over the ^13^C_3 15_N_1_-labeled internal standard was integrated and interpolated against a calibration curve that was linear over the range of the samples. Protein abundance from the extracted pellet was measured by BCA assay (Thermo Fisher).

### Acid Extraction of histones for quantitative proteomics

Histones were extracted according to Schecter et al. with modifications^6^. Briefly, 1.5 × 10^6^ cells were plated for seeding overnight. The following day, cells were treated accordingly for 24 h. Media was then removed, and cells were washed once, scraped into ice-cold PBS (1 mL), and pelleted via centrifugation (1,000 × *g*, 5 min, 4°C). Pellets were lysed in hypotonic lysis buffer (10 mM HEPES/KOH pH = 7.9, 1.5 mM MgCl_2_, 10 mM KCl, 5 mM sodium butyrate (NaB), 0.5% IgePal, and protease and phosphatase inhibitor cocktails at 1:100 v/v [Sigma Aldrich]) for 1h on ice and nuclei were pelleted via centrifugation (250 × *g*, 30 min, 4°C). Pellets were resuspended in 200 µL 0.4N H_2_SO_4_ and suspensions were rotated end-over-end at 4°C for 3 h. Non-histone proteins were pelleted via centrifugation at 1000 × *g* for 10 min at 4°C. Soluble histones (supernatant) were transferred to a new tube and precipitated via the addition of 50 µL 100% TCA (w/v) yielding a final concentration of 20% TCA. Suspensions were incubated on ice for 1h with occasional vortexing. Histones were pelleted via centrifugation at 16,000 × *g* for 10 min at 4°C and the supernatant was carefully removed with a pipette tip. Histones are seen as a ‘smear’ on the side of the tube and a small pellet. Histones were then washed with acidified acetone (0.1% (v/v) HCl) followed by two washes with ice-cold acetone. Histones were allowed to air-dry to remove any residual acetone. For western blotting, samples were resuspended in 200 µL ddH_2_O.

### Quantitation of histone peptides

Dried histones (above) were resuspended in 50 µL of 50 mM NH_4_HCO_3_, 5 µL of 1 M NH_4_OH and 100 µL propionic anhydride (prepared as a 1:3 v/v in isopropanol). The pH was adjusted to 8.0 and samples were incubated at 52°C for 1 h. Samples were then dried to completion in a SpeedVac and resuspended in 100 µL 50 mM NH_4_HCO_3_, containing 1 µg of sequencing-grade trypsin (Promega). Samples were digested overnight at 37°C and then subjected to an additional round of propionylation via the addition of 100 µL propionic anhydride solution (1:3 v/v isopropanol) and 25 µL NH_4_OH to maintain a pH of 8.0. Samples were incubated at 52°C for 1 h and dried to completion in a SpeedVac. Dried histones were resuspended in 200 µL of 0.1% formic acid in H_2_O (i.e. buffer A) and subjected to MRM-MS.

12 µL of peptide suspensions were chromatographed using a Shimadzu LC40 system equipped with a 2.1 mm × 100 mm, 1.8 µm particle diameter Acquity UPLC HSS C_18_ Column (Waters, Milford, MA) at a flow rate of 0.4 mL/min. Buffer A (0.1% formic acid in ddH_2_O) and Buffer B (0.1% formic acid in ACN) were used with the following gradient: 1 min, 1% B; 20 min, 35% B; 24 min, 98% B; 27 min, 98% B; 27.7 min, 1% B. The column was equilibrated at 1% B for 2 min between runs. MRM was performed in positive ion mode using an AB SCIEX 6500+ QTRAP with the parameters defined in **Table S6**. A minimum of 3 transitions were used for each peptide. Peak areas were calculated and summed with all peaks of the same peptide. Modified peptides were then divided by this sum, providing a relative percent abundance of the modification at that site.

### Isolation of nuclei from macrophages for ATAC-seq

1 × 10^6^ cells were plated and treated as described above. The following buffers were prepared prior to nuclear isolation: Omni Resuspension Buffer^7^ (RSB, 10 mM Tris-HCl pH 7.4, 10 mM NaCl, 3 mM MgCl_2_, stored at 4°C), 2x Omni TD Buffer (20 mM Tris-HCl pH 7.6, 10 mM MgCl_2_, 20% dimethyl formamide, stored at - 20°C), freezing buffer^8^ (50 mM Tris-HCl pH 8.0, 5 mM Mg(OAc)_2_, 25% glycerol, 0.1 mM EDTA, stored at -20°C). Following treatments, media was removed, and cells were washed with 1 mL ice-cold PBS. Cells were then scraped into 1 mL of ice-cold PBS and pelleted (1,000 × *g*, 5 min, 4°C). Supernatant was removed and lysis buffer (250 µL, 1X resuspension buffer, 0.1% Igepal-CA630, 0.01% digitonin, 0.1% Tween-20) was added to each sample. After incubation on ice (3 min), 1 mL of stop solution (1X resuspension buffer, 0.1% Tween-20) was added. Nuclei were then counted (10 µL nuclei, 40 µL of 2x Omni TD, 50 µL trypan blue). Samples were once again pelleted (500 × *g*, 10 min, 4°C), and resuspended in freezing buffer (1 mL, 1X freezing buffer, 5 mM dithiothreitol, 2% protease inhibitors (v/v)). Samples were flash frozen and analyzed accordingly.

### ATAC-seq library preparation

ATAC-seq was performed using the Omni-ATAC protocol with slight modifications^7^. The flash-frozen nuclei isolated from RAW264.7 macrophages were removed from the liquid nitrogen and thawed in a water bath at 37°C for 1 to 2 min until only a tiny ice crystal remained. After thawing, the nuclei were diluted with 3 mL ATAC resuspension buffer (RSB) supplemented with 0.1% Tween-20 and 0.1% BSA (RSB washing buffer) and centrifuged at 500 r.c.f for 10 min in a pre-chilled (4°C) swinging-bucket centrifuge. The RSB buffer was prepared to consist of 10 mM Tris-HCl (pH 7.5, Invitrogen, cat. no. 15567027), 10 mM NaCl (Invitrogen, cat. no. AM9759), and 3 mM MgCl_2_ (Invitrogen, cat. no. AM9530G) in nuclease-free water. The nuclei pellet was resuspended with another 1 mL of RSB washing buffer and transferred to a 1.5 mL LoBind tube followed by centrifugation at 500 r.c.f for 5 min in a pre-chilled fixed-angle centrifuge. After pelleting, nuclei were resuspended in 100 µL of PBS containing 0.04% BSA (PBSB), counted, and diluted to 3,030 nuclei/µL with PBSB. Because the nuclei are sensitive to osmotic pressure and will inflate in Trypan blue solution, nuclei in PBSB 1:1 to 2X Omni tagmentation buffer was added (containing 20 mM Tris HCl pH 7.5, 10 mM MgCl2 and 20% Dimethyl Formamide) followed by adding 1 volume of Trypan blue for counting. A volume of 6.6 μL of diluted nuclei (20,000 total nuclei) was transferred to 12.4 μL of transposition reaction (10 μL of 2X Omni tagmentation buffer, 0.2 μL of 1% digitonin, 0.2 μL of 10% Tween-20, and 2 μL of H_2_O). Then, 1 μL of 300 μg/mL Tn5 transposase^9^ to the transposition mix containing nuclei was added and performed tagmentation on a thermocycler at 37°C for 30 min. The tagmented DNA was cleaned up using Zymo DNA Clean and Concentrator-5 columns with 5X binding buffer (Zymo, cat. no. D4004). 5 μL of purified DNA out of 12.5 μL purified product was amplified in a 25 μL PCR reaction containing NEBNext PCR master mix (1X final), 0.5X SYBR Green and 1.25 μM of Ad1 primer from PMID: 24097267^10^ and 1.25 μM of an N7 primer containing custom barcodes (**Table S5**). Samples were amplified on a Bio-Rad CFX Connect Real-time cycler using the following program: 72°C for 5 min; 98°C for 30 s; 11 cycles of 98°C for 10 s, 63°C for 30 s, 72°C for 1 min. PCR products were cleaned up with Ampure XP beads. A double size selection was performed to remove DNA fragments larger than ∼1500 bp (with 0.4X AMPure XP beads) and smaller than ∼100 bp (with 1.5X AMPure XP beads). To do this, we added 25 μL of Qiagen Buffer EB to each PCR reaction to bring the volume to 50 μL, and then added 20 μL of beads (homogenizing the mixture well by pipetting), followed by a 5 min incubation at RT. The samples were then put on a magnet stand to bind the beads. 68 μL of the supernatant was transferred to a new 1.5 mL tube and an additional 53 μL of beads were added to the supernatant and resuspended thoroughly (we estimated that 19.4 μL of the 68 μL transferred supernatant was bead buffer and 48.6 μL was sample, so to get to 1.5X beads for the second selection we needed to add an additional (72.9 - 19.4) μL of beads, which we rounded to 53 μL). After another 5 min at RT, the samples were placed on the magnet to clear the beads and the supernatant was discarded. Beads were washed twice with 200 μL of 80% EtOH (made fresh for each experiment). After the second wash, the beads were briefly spun, put back on the magnet stand, and residual ethanol was removed. The beads were then air dried for 1 min. Beads were then removed from the magnet and resuspended in 22 μL of Buffer EB, incubated for 2 min at RT and then placed on the magnet stand again. 20 μL of supernatant was transferred to a new tube. Following this Ampure size selection, the ATAC-seq libraries were quantified by Qubit 1X dsDNA HS Assay Kit and run on a 6% PAGE gel prior to sequencing.

### Sequencing

All ATAC-seq libraries were pooled together and sequenced on a NextSeq 550 Platform with 51 bp for each read.

### ATAC-Seq data analysis

The specific programs (and their version) used in data analysis were as follows: Trimmomatic v0.36^11^, SAMtools v1.4^12^, Picard v2.20.2 [Picard toolkit. Broad Institute, GitHub repository. 2019. Available: http://broadinstitute.github.io/picard/], Bowtie2 v2.2.9^13^, MACS2 v2.1.2^14^, bedtools v2.28.0^15^, deepTools v3.5.1^16^, R v4.1.1 [R Core Team. R: A Language and Environment for Statistical Computing. 2021. Available: https://www.R-project.org/], edgeR v3.40.0^17^, rGREAT (v2.0.2)^18^ and KEGGREST package [CITE: Tenenbaum D, Maintainer B (2023). KEGGREST: Client-side REST access to the Kyoto Encyclopedia of Genes and Genomes (KEGG). R package version 1.40.0.]

### Preprocessing

The paired-end reads were preprocessed using trimmomatic to trim the Nextera adaptors and low quality reads with the parameter setting as “LEADING:3 TRAILING:3 SLIDINGWINDOW:4:10 MINLEN:20”. The trimmed reads were then mapped to mm10 mouse genome reference using Bowtie2. The parameters “-X 2000” and “-3 1” were used to restrict the maximum fragment length of 2000 bp and trim 1 base from the 3’ end of each read before alignment. Following mapping, we retained the reads that confidently mapped to the assembled nuclear chromosomes (MAPQ ≥ 10) and were in proper pairs (specified by the “-f3” and “-F12” options in SAMtools), and then removed duplicate reads using Picard “MarkDuplicates” tool.

### Peak calling

To call peaks on each condition, we combined the deduplicated reads for all replicates from the same condition and then identified peaks using MACS2 by considering a 200 bp window centered on the read start using the parameters “--nomodel --keep-dup all --extsize 200 --shift -100”. Because each peak may have multiple summits (and will therefore be listed multiple times in the resulting peak bed file), the peaks output from MACS2 were then merged into a single peak set for each sample using bedtools “merge”. Any peaks overlapping with the problematic regions defined by ENCODE blacklist^19^ were removed.

### Differential analysis

A peak-count matrix was generated using deepTools “multiBamSummary” with a common peak set combining all peaks across conditions as rows and samples as columns. The edgeR package was used to identify differentially accessible peaks between each comparison. In brief, the lowly accessible peaks were filtered out using the edgeR “filterByExpr” function with default parameters followed by calculating the normalization factors using the trimmed mean of M-values method through the “calcNormFactors” function. Then, we estimated dispersions using “estimateDisp()” with a design matrix and “robust = TRUE” and performed hypothesis testing using the quasi-likelihood F-test implemented in edgeR. The Benjamini and Hochberg (BH) method^20^ was used to control the false discovery rate (FDR). A two-threshold approach^21^ was used to determine the peaks shared between conditions by requiring a stringent 5% FDR threshold for at least one test, but allowing for a more relaxed threshold (20% FDR) for the other test.

### Functional analysis

The KEGG pathways enrichment analysis was performed by applying rGREAT on differential peaks using a binomial test. The gene sets of KEGG pathways were retrieved using the KEGGREST package.

### Statistical Analysis

All statistical tests were performed in GraphPad Prism 10.0. The specific statistical analysis is mentioned in each figure legend. For two-way ANOVA: all tests were performed with two-tailed analysis. Error bars are reported as standard deviation of the mean (SD) standard error of the mean (SEM), specified in the figure legends. All N values are provided in the figure legends.

